# Two mechanisms of photoenergy regulation revealed by kinetic behaviors of chlorophyll fluorescence during light adaptation

**DOI:** 10.1101/2024.07.14.603445

**Authors:** Junqing Chen, Lijiang Fu, Ya Guo, Jinglu Tan

## Abstract

Photochemical reactions were analyzed and modeled to observe photoenergy regulation mechanisms in PSII from measured chlorophyll fluorescence (ChlF) kinetics. Two pH-driven mechanisms were revealed: One is the PsbS-mediated non-photochemical quenching (NPQ) of activated antennae (A*), as commonly understood, and the other is a disruption of energy transfer from A* to P680. Representing the latter with a conformational change from complex formation of zeaxanthin and lutein with antennae closely described measured ChlF from initially dark-adapted state to light-adapted state and measured NPQ variations. Analysis based on the model indicates that zeaxanthin and lutein lead to slow protracted reductions in P680* and ChlF via the second mechanism without strong influence by pH or PsbS. Protonation of PsbS plays a major role via the first mechanism in responding to fast changes in illumination. The research provides insights into the mechanisms for photoenergy regulation in PSII and a kinetic model with broadened applicability.

## Introduction

As commonly understood, photons captured by plant photosystem II (PSII) antennae have three possible fates: photochemical reactions (photosynthesis), non-photochemical quenching (NPQ), and emission as chlorophyll fluorescence (ChlF)(Matuszy, 2016). Photochemical reactions accumulate protons in the thylakoid lumen to rotate the ATP synthase by transporting energized electrons originating from separation of water molecules(Junge and Nelson, 2015). Under strong illumination, photoprotection is activated to minimize the harmful effects of excess light(Anderson, Park and Chow, 1997). NPQ is considered the primary photoprotection process, which releases excess absorbed photoenergy as heat. NPQ is known to include qE (quenching of energized electrons), qT (state transitions), qI (photoinhibition)(Lambrev *et al*., 2012; Barbara *et al*. 2014; Nagao *et al*., 2019), qZ (zeaxanthin-dependent quenching), and qH (sustained quenching)(Bassi and Dall’Osto, 2021). The proton gradient across the thylakoid membrane triggers qE(Hager and Holocher, 1994; Goss, Richter and Wild, 1995) by activating the de-epoxidation of violaxanthin to form zeaxanthin (Zea)(Färber *et al*., 1997) and lutein (Lut)(Bungard *et al*., 1999), reactions known as the reversible enzymatic xanthophyll cycle and lutein epoxide cycle(García-Plazaola, Matsubara and Osmond, 2007), respectively. The protonation of the PsbS protein is believed to activate quenching sites in antennae(Barbara *et al*. 2014; Pinnola and Griffiths, 2019). qT is the process to transfer phosphorylated light-harvesting proteins from PSII to PSI(Barbara *et al*. 2014). qI permanently degrades PSII(Bassi and Dall’Osto, 2021). Furthermore, qZ was observed after eight minutes of photosynthesis(Bassi and Dall’Osto, 2021)and depends on Zea but not on PsbS and lumen pH(Nilkens *et al*., 2010). qH is ahead of qI and reversibly decreases the function of PSII. It is activated by a plastidial lipocalin (LCNP), which is regulated by a quenching suppressor SOQ1(Malnoë *et al*., 2018). In addition, antennae were found to avoid absorbing photons and the concentration of excited sites is reduced under excess light, which is referred to as qM(Cazzaniga *et al*., 2013).

While the phenomena and roles of NPQ are well established, the mechanisms underlying the interaction of carotenoids with antennae to regulate photoenergy transfer remain less clear(Bassi and Dall’Osto, 2021). Two prevailing hypotheses are: (1) Binding of zeaxanthin(Zea)/lutein(Lut) with LHCII may lead to a conformational change to channel energy to carotenoid quenchers(Dall’Osto *et al*., 2017; Bassi and Dall’Osto, 2021), and (2) Excited chlorophylls (Chl) may transfer energy though a “charge-transfer state” by a Chl-Zea heterodimer, typically observed in monomer LHCs(Bassi and Dall’Osto, 2021). qE has an impact on Zea^+^ formation(Holt *et al*., 2005); besides, both Zea and Lut have been shown to contribute to the quenching activity of monomers(Dall’Osto *et al*., 2017). These quenching processes are contingent upon the presence of PsbS(Dall’Osto *et al*., 2017). The absence of monomers (CP29, CP26, and CP24) results in the peripheral LCHII “disconnecting” from the PSII core and causes an over-accumulation of trimers(Dall’Osto *et al*., 2014, 2017), signifying a role of monomers as a “bridge”(Dall’Osto *et al*., 2020). During NPQ, monomers may function as a modulator in the quenching process(Dall’Osto *et al*., 2017) and PsbS may participate in the dissociation of CP24 and CP29 complexes, which serve as bridges between the internal and external antenna layers(Betterle *et al*., 2009).

Modeling is a powerful tool for revealing the underlying processes or states of a complex system based on observable variables. ChlF has been extensively used as such an observable variable. Because ChlF energetically complements and depends on NPQ and photochemical reactions, a mechanistically based model would provide further insights into the NPQ process from ChlF measurements. Many models have been developed for PSII phototransduction kinetics(Van Kooten, Snel and Vredenberg, 1986; Guo and Tan, 2011), light-harvesting antenna(Amarnath *et al*., 2016), ion flux and proton motive force across the thylakoid membrane(Davis *et al*., 2017), qE process(Zaks *et al*., 2012), and Zea- and Lut-dependent NPQ(Leuenberger *et al*., 2017). Van Kooten et al. propsed a model of phototransduction to provide a brief structure of photosynthesis(Van Kooten, Snel and Vredenberg, 1986). Other models were developed with a specific focus such as energy transfer by the intermediates based on observed ChlF(Zhu *et al*., 2005; Guo and Tan, 2011). These models nearly always focus on the fast ChlF induction commonly referred to as the OJIP phases. Since the NPQ process was discovered about 50 years ago, models of quenching have been created. Ebenhöh et al. presented a simple mathematical model of NPQ and analyzed the rate constants in the quenching dynamics(Ebenhöh *et al*., 2011). Zaks et al. developed a quantitative model to describe and analyze the qE component(Zaks *et al*., 2012). They used a set of equations to represent the PsbS function in activating the quenching sites(Bennett *et al*., 2019) and stated that the Zea concentration determined the maximum of qE(Zaks *et al*., 2012), but the model was not completely consistent with experimental data when the light-adaptation process progressed. Leuenberger et al. discussed the biochemical quenching mechanism by devising a model for Zea- and Lut-dependent quenching based on fluorescence lifetime measurements(Leuenberger *et al*., 2017). Rarely has a model represented the ChlF kinetics of both the OJIP and the light-adaptation phases with a good level of accuracy. An opportunity, therefore, exists to develop a model that would allow observation of the photoenergy regulation processes from ChlF measurements.

In this research, we aimed to gain an understanding of the mechanisms of photoenergy regulation by synthesizing the current knowledge in PSII photoenergy transduction and quenching through the development of a model for ChlF kinetics from dark-adapted state to light-adapted state. The broadened model applicability required a detailed analysis of the major reactions involved. The model was evaluated against measured ChlF and compared with published NPQ data. The analysis and experiments reveal two pH-mediated mechanisms for PSII photoenergy regulation and provide important insights into how plants respond to lighting variations and protect themselves from photodamage.

## Model Summary

Figure 1 broadly illustrates the PSII reactions, variables, and processes of photoelectron transport and NPQ analyzed to develop a kinetic model. The model consists of differential equations for 14 variables of reaction center or thylakoid lumen conditions (*A*^∗^, *P*680^∗^, 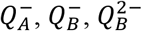, *PQ, H*^+^ *pH, K*^+^, *Cl*^−^, *Anth, Zea, Lut*, and *A*_*c*_), a controllable input (illumination intensity *u*), and an observable or measurable output (*ChlF*). The model has 30 parameters to be estimated from measured ChlF as detailed in the Methods section. A complete description of the analysis and model is provided in the Supplementary Materials. The NPQ reactions involving *H*^+^, *pH, K*^+^, *Cl*^−^, *Anth, Zea, Lut*, and *A*_*c*_ are analyzed and modeled as discussed below.

**Fig. 1.**
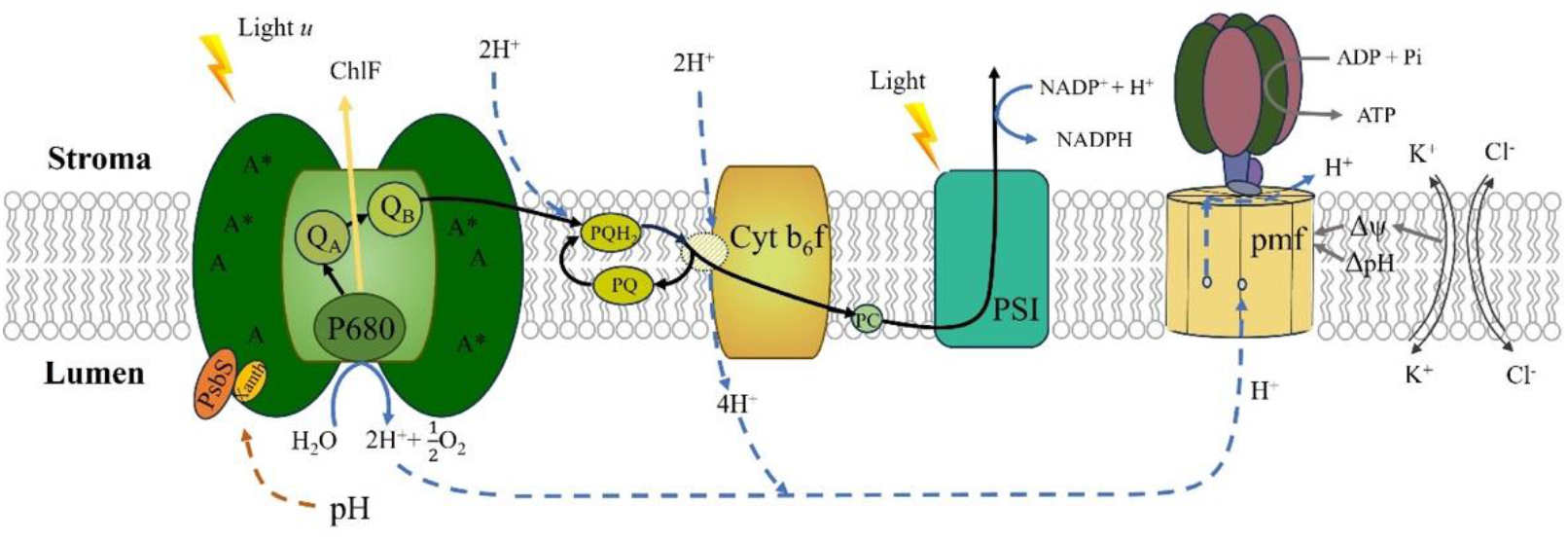
An illustration of the main processes and variables modeled plus those of the immediate downstream. After a PSII antenna (A) is activated by a photon to become A*, the energy will transfer among the chlorophylls in the antennae. When a P680 captures the energy to become the excited form, P680*, it may pass its energized electron via a pheophytin to plastoquinone A and B (Q_A_ and Q_B_) and then to plastoquinone (PQ) and receive a replacement electron originating from water, or it may relax to release chlorophyll fluorescence (ChlF). PQ undergoes a redox reaction with two protons (H^+^) from the thylakoid stroma, forming PQH2. Interaction with cytochrome b6f (Cyt b6f) engages two additional protons, resulting in the release of four protons into the lumen. Detailed analysis and modeling of these reactions are included in the Supplementary Materials. H^+^ accumulation in the lumen changes the pH difference across the membrane (ΔpH) and the membrane potential (Δ*ψ*) formed by H^+^ and other ions (represented by K^+^ and Cl^-^). This results in a proton motive force (pmf) to drive subsequently ATP synthesis. Low lumen pH also initiates de-epoxidation of xanthophylls into antheraxanthin (Anth), zeaxanthin (Zea), or lutein (Lut), which are catalyzed by protonated PsbS to quench A* or to alter photoenergy transfer as detailed in the main text.

### Ion fluxes and pH

#### Ion fluxes

As commonly known (details in Supplementary Materials), photochemical reactions produce a net increase in H^+^ concentration in the thylakoid lumen, which changes the electric potential difference (Δ*ψ*) and pH difference (ΔpH) across the thylakoid membrane. These differences form a proton motive force (pmf) that drives ion fluxes (H^+^ and others). Δ*ψ* is determined from the charges of cations and anions on both sides of the membrane by Coulomb’s law. The major cations and anions are potassium (K^+^) and chloride (Cl^-^), which may be used to account for the effects of all ions. Δ*ψ* and the rates of proton and ion fluxes from the lumen to the stroma can be represented as:

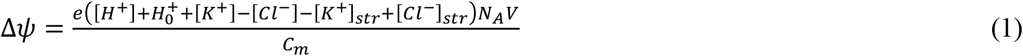

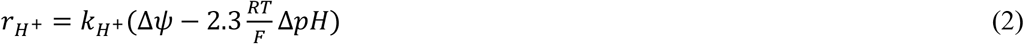

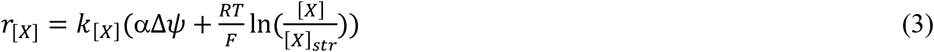

where *e* is the elementary charge, 1.602176634’10^−19^ C; *N*_*A*_ is Avogadro’s number, 6.022’10^23^; and *V* is the thylakoid lumen volume, 2.1’10^−15^L (Beebo *et al*., 2013); *C*_*m*_ is electrical capacitance of the thylakoid membrane, 0.6 mF/cm^2^ (Davis *et al*., 2017); [*x*] is the molar concentration of ion species *x* in the lumen; subscript *str* signifies stroma; [*H*^+^] is the net change in total proton concentration (both bond and free) from the initial dark-adapted condition or the total net production of protons in the lumen; and *H*^+^ is initial proton concentration *difference* between lumen and stroma (Since the stroma conditions are assumed unchanged, [*H*^+^] is also the change from the initial proton concentration difference *H*^+^); *R* is the gas constant, 8.3145 J/mol/K; *T* is absolute temperature in K; *F* is the Faraday constant, 96,485 C/mol; Δ*pH* is the pH difference between the lumen and the stroma (Δ*pH* = *pH-pH*_*str*_) with the pH in stroma *pH*_*str*_ set to 8 (Werdan, Heldt and Milovancev, 1975); *K*_[*X*]_ is a constant accounting for the conductivity or permeability of ion species x through the thylakoid membrane; α = 1 for cation and -1 for anion. The initial conditions are discussed in the Methods section.

#### Buffer capacity

pH in the thylakoid lumen depends on the buffer capacity and proton concentration. There are multiple buffering species in the lumen. The overall buffering species concentration was reported to be 300 mM (Van Kooten, Snel and Vredenberg, 1986) and the overall acid dissociation constant in the lumen was reported to be approximately 5.5 (Heldt *et al*., 1973). Therefore, instead of assuming a constant buffer capacity of 0.03 as often done in prior work, the buffer capacity in the lumen can be expressed as a function of pH as:

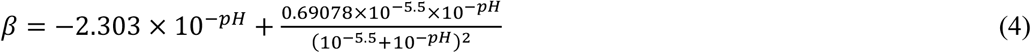

### Non-photochemical quenching

Non-photochemical quenching (NPQ) refers to self-protection mechanisms to divert excess absorbed light energy. Xanthophylls, Photosystem II subunit S protein (PsbS), and antennae are known to be the necessary components (Niyogi, 2000).

#### Activation of enzymes and PsbS

Violaxanthin de-epoxidase (VDE) activated by low lumen pH is required in the epoxide cycle of both violaxanthin and lutein(García-Plazaola, Matsubara and Osmond, 2007). Zeaxanthin epoxidase (ZE), located in the stroma, catalyzes the reverse reaction of xanthophyll(Latowski and Grzyb, 2004). The lumen pH is also known to activate PsbS, leading to a conformational change in LHCII to enable quenching of activated antenna A* by pigments. These enzymes and PsbS are large molecules compared with proton(Pinnola and Griffiths, 2019); therefore, the proportion of protonated VDE, ZE, and PsbS can be expressed by the Hill equation

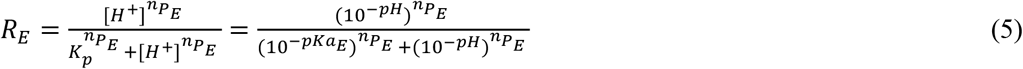

where *E* represents VDE, ZE, or PsbS; and *pKa*_*E*_ and 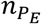 are the pKa and the Hill coefficient, respectively, for protonation of *E*.

#### Epoxide cycle of xanthophylls

The xanthophyll cycle was first discovered in 1962 through the successful extraction of antheraxanthin (Anth) and zeaxanthin (Zea) (Yamamoto, 1962). Low thylakoid lumen pH activates VDE on the lumen side of the membrane by protonation and activated VDE catalyzes the de-epoxidation of violaxanthin (Vio) to Zea via intermediate Anth, while ZE converts the intermediates back to Vio(Latowski and Grzyb, 2004). The VDE enzyme may also be able to convert lutein epoxide (Lx) to lutein (Lut) (García-Plazaola, Matsubara and Osmond, 2007). Based on the Michaelis-Menten enzymatic kinetics, the net forward epoxidation rates are

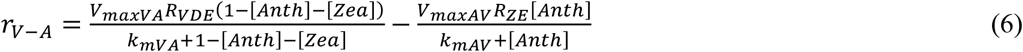

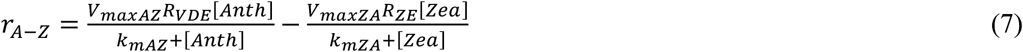

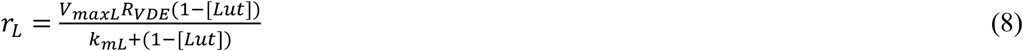

where *V*_*maxVA*_ is the maximum rate of conversion from Vio to Anth; *K*_*maxVA*_ is the Michaelis constant of the reaction from Vio to Anth; *V*_*maxAZ*_ and *K*_*maxAZ*_ are, respectively, the maximum rate and Michaelis constant of the reaction from Anth to Zea; *V*_*maxAV*_ and *K*_*maxAV*_ are, respectively, the maximum rate and Michaelis constant of the reaction from Anth to Vio; *V*_*maxZA*_ and *K*_*maxZA*_ are, respectively, the maximum rate and Michaelis constant of the reaction from Zea to Anth; *V*_*maxL*_ and *K*_*mL*_ are, respectively, the maximum rate and Michaelis constant of lutein formation; [*Anth*], [*Zea*], and [*Lut*] are, respectively, the probabilities or fractions of Vio in the form of Anth or Zea, and Lx in the form of Lut, which implies that the sum of [*Vio*], [*Anth*], and [*Zea*] is 1 (or 100%) and the sum of [*Lx*] and [*Lut*] is 1; and finally, *R*_*ZE*_ is the proportion of protonated ZE, which is negligible under high illuminations. The use of fraction instead of concentration as unit here saves the need for determining the initial concentrations of Vio and Lx, which are included in the *V*_*max*_ and *K*_*m*_ values.

#### Photoenergy quenching and transfer

The current prevailing view is that an incident light is absorbed at a certain rate to produce the excited form of antennae A*, which have three destinations: photochemical reactions, fluorescence generation, and heat production (or NPQ). Each A* has a chance to go to one and only one of the three destinations; i.e., the three compete for the same A* population. As the exact molecular mechanism is not yet clear, the rate of A* quenching by an NPQ carotenoid may be considered, for simplicity, as a first-order concentration-driven reaction mediated by *R*_PsbS_. The total rate of quenching by Lut, Zea and Anth can thus be written as

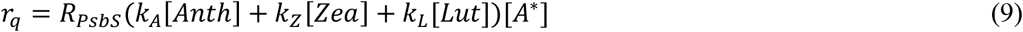

where *K*_*X*_(*X* = *A, Z* or *L*) are the reaction (quenching) rates for Anth, Zea and Lut, respectively. The initial concentration (number of molecules per reaction center) of Lx is lumped into *K*_*L*_ and that of Vio into *K*_*A*_ and *K*_*Z*_. This mechanism of pH-mediated NPQ is reversible when pH and thus *R*_PsbS_ change.

The protonation of PsbS also leads to Zea- and Lut-dependent interactions with the monomeric LHC protein(Dall’Osto *et al*., 2017; Pinnola and Griffiths, 2019), changing the behavior of the monomeric LHC. Researchers have observed that the concentration of functioning antennae appears to be reduced under strong lighting and characterized this reduction as antenna avoidance(Cazzaniga *et al*., 2013) or absorption area reduction(Cazzaniga *et al*., 2013). The exact mechanism is yet to be determined, but some suggest that activated PsbS may catalyze the formation of antenna-carotenoid complexes(Pinnola *et al*., 2015). Besides, sharp declines in PSII photochemistry rate have been observed during light adaptation(Pfündel *et al*., 2013). The complexed antennae are thus effectively disabled for photoenergy transfer as observed by Erhard et al. (Pfündel *et al*., 2013) and Luca et al.(Dall’Osto *et al*., 2017) Also, this effect is only slowly reversible(Dall’Osto *et al*., 2017). Monomers are believed to be the primary energy transfer pathway between A* and P680 (Dall’Osto *et al*., 2014; Nicol, Nawrocki and Croce, 2019).

We use the suggested complex formation to represent the Zea- and Lut-dependent antenna conformational changes that disrupt photoenergy transfer from antennae to P680. With protonated PsbS as enzyme, the Michaelis-Menten equation can be used to describe the rate of antenna-carotenoid complex (A_c_) formation from Zea and Lut, respectively:

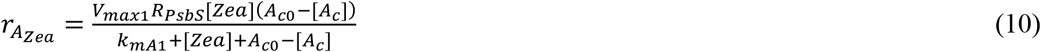

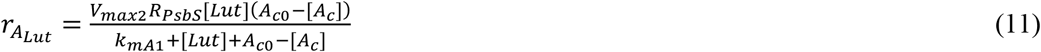

where *V*_*max*1_ and *K*_*maxA*1_ are the maximum rate and Michaelis constant of A_c_ formation from Zea, respectively; *V*_*max*2_ and *K*_*maxA*2_ are the maximum rate and Michaelis constant of A_c_ formation from Lut, respectively; [*A*_*c*_] is the fraction of antennae formed into complex; and *A*_*c*0_ is the maximum fraction of antennae (both monomers and trimers) that can form A_c_, which is assumed to be 0.85 (Wentworth, Ruban and Horton, 2004);

A_c_ complex formation leads to a reduction in the number of A* able to transfer energy to P680 by the proportion of complexed antennae, [*A*_*c*_], as shown in Eqn. (12),

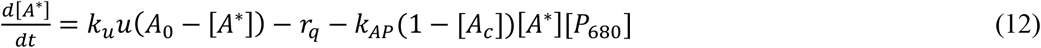

where *K*_*u*_ is the energy absorption rate from photons; *u* is illumination intensity; *A*_0_ is the number of antennae in one reaction center (set to 290 (Guo and Tan, 2011; Junge and Nelson, 2015)); *r*_*q*_ is the A* quenching rate by carotenoids (Eqn. 9); *K*_*AP*_ is the rate of energy transfer from activated antenna (A*) to P680; and [*x*] represents the number or fraction of species *x*. Eqn. (12) means that the time rate of change for A* consists of one production term (by illumination *u*) and two consumption terms: quenching by carotenoids and transfer of energized electrons to P680 modulated by A_c_.

In summary, the pH-mediated regulation of photoenergy transfer (which may be generally referred to as NPQ activities) occurs via *two separate mechanisms*:

1. Quenching of A* by carotenoids (Eqn. 9), and
2. Antenna-carotenoid complex (A_c_) formation disabling transfer of energy from A* to P680 (Eqns. 10 and 11)

Mechanism 1 represents what is commonly referred to as the qE process (quenching of A*) while mechanism 2 represents disconnection of energy transfer pathway from A* to P680 rather than quenching *per se*. These two mechanisms will be further illustrated and discussed in later sections.

#### Other NPQ components

There are other significant NPQ components; namely, qT, qH, qM, and qI. qT is a component caused by the movement of phosphorylated LHCII antennae from PSII to PSI. This reversible reaction usually happens in low-light conditions(Barbara *et al*., 2014). The measurement of ChlF was done in this work under the conditions of actinic light and pulses of high intensities and thus qT would not be present. qH is a process that happens earlier than qI (and thus the name)(Malnoë, 2018).

Both qI and qH decrease the light adsorption, but qH is reversible. qH is catalyzed by a plastidial lipocalin (LCNP), which decreases the light absorption by antennae. Zea, PsbS, and Lut have been shown to be unrelated to the qH process. The exact mechanism is still unknown, but it is well possible that the underlying NPQ mechanisms of qH is similar to those of the qE process(Malnoë, 2018). It has been stated that a light intensity of 1,200 umol/m^2^/s would not activate LCNP(Malnoë *et al*., 2018). Experiments conducted in this research with an illumination of 1,000 umol/m^2^/s should thus not include significant effects of the qH process. qM was found to cause a decline in ChlF because of decreased light absorption after 20 minutes(Cazzaniga *et al*., 2013). If the qH and qM effects were actually significant for the experiments conducted in this work, they could be considered as accounted for in the two mechanisms.

*y* is the measured ChlF, n is the number of the data points, and ŷ is the model-predicated ChlF.

## Results

### Model validation and analysis

Figure 2 shows the model prediction of ChlF compared with measurement for a spinach sample. The curves may be viewed in three segments: the commonly called OJIP phases from the initial dark-adapted state till about 1 s shown with an inset in log scale, a light-adaptation period of ChlF decline often referred to as the PSMT phases, and the responses to illumination pulses after 300 s shown with another magnified inset. Figures for additional samples are included in Fig. S1 in Supplementary Materials.

**Fig. 2.**
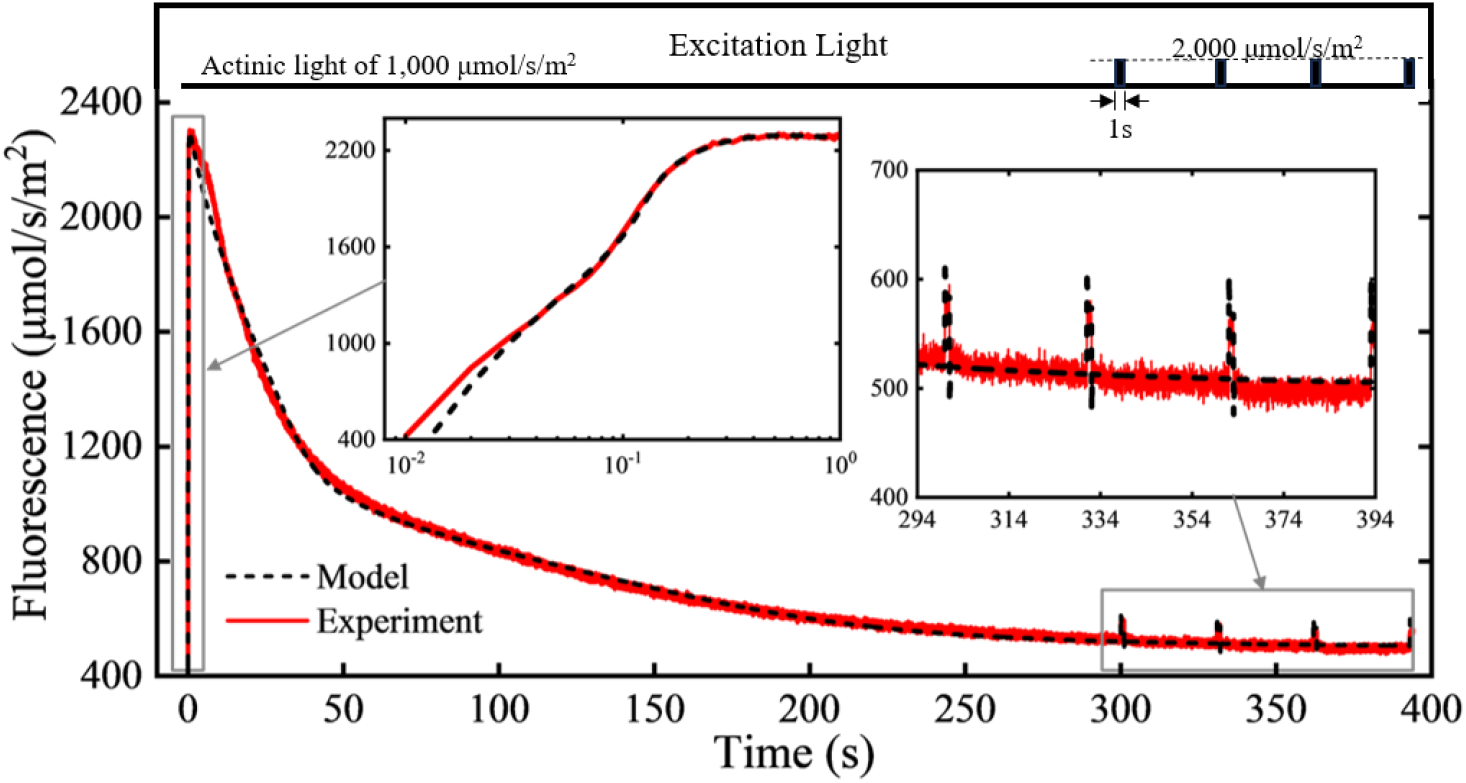
Comparison of experimental ChlF measurements and model predictions. The sample was an initially dark-adapted spinach leaf in 22°C. The illumination was a 1,000 mmol/s/m^2^ constant actinic light starting at time 0 with 1-s wide pulses of 1,000 mmol/s/m^2^ above the constant light. The light source was the built-in white LED of an OS5p+ Chlorophyll Fluorometer (Opti-Sciences, Hudson, NH, USA). One inset shows the OJIP phases in the first second in log scale and another shows a magnified view of the pulse responses. Figures for additional samples are included in Supplementary Materials.

The model predictions and experimental measurements are in good agreement based on the MAPE and R^2^ values. The 95% confidence interval of the MAPE for all the samples is [1.962%, 3.512%]. The low MAPE (defined in Methods) values represent small percentage errors and close representation of the measured ChlF by the model. The 95% confidence interval of R^2^ is [0.989, 0.997], again indicating that the model closely follows the measured ChlF behavior.

It should be noted that the model is able to represent measured ChlF in all three segments. As the log-scale inset shows, the model follows the OJIP phases till the peak at about 1 s, which is the commonly used ChlF induction curve measured after dark adaptation. The model also follows the nonlinear behaviors in the PSMT phases, which show a rapid decline from the peak at about 1 s till about 50 s, a decreased rate of decline till about 250 s, and further leveling off after that. In response to the pulses of 2,000 mmol/s/m^2^ (1,000 mmol/s/m^2^ above the constant actinic light of 1,000 mmol/s/m^2^), the amplitude of ChlF rise is obviously lower than the peak around 1 s because of light adaptation, as is commonly known. It is important to note that, as shown in the inset, the model closely represents the amplitude of the pulse responses. This is significant because such pulse responses are commonly used to measure NPQ and photosynthetic activities under light-adapted conditions.

### Kinetic variations

Figure 3 shows the modeled dynamic behaviors of 21 variables as functions of time with the model parameter values obtained from the sample shown in Fig. 2. With the onset of lighting, A* and P680* increase, and the reduced Q_A_ and the double-reduced Q_B_ rapidly increase during the OJIP phases while the single-reduced Q_B_ is the intermediate balancing the two. This is accompanied by a rapid decrease in PQ and increase in H^+^, resulting in a rapid pH drop in the thylakoid lumen to below 6 in about 1 s. The low pH activates VDE, initiating the production of Auth, Zea and Lut; and also protonates PsbS to trigger the PsbS-mediated quenching of A* (qE) by these catenoids at the rate of *r*_*q*_ (mechanism 1). *r*_*q*_ increases in the early part of the PSMT phases and remains high during light adaptation because of elevated Zea and Lut.

**Fig. 3.**
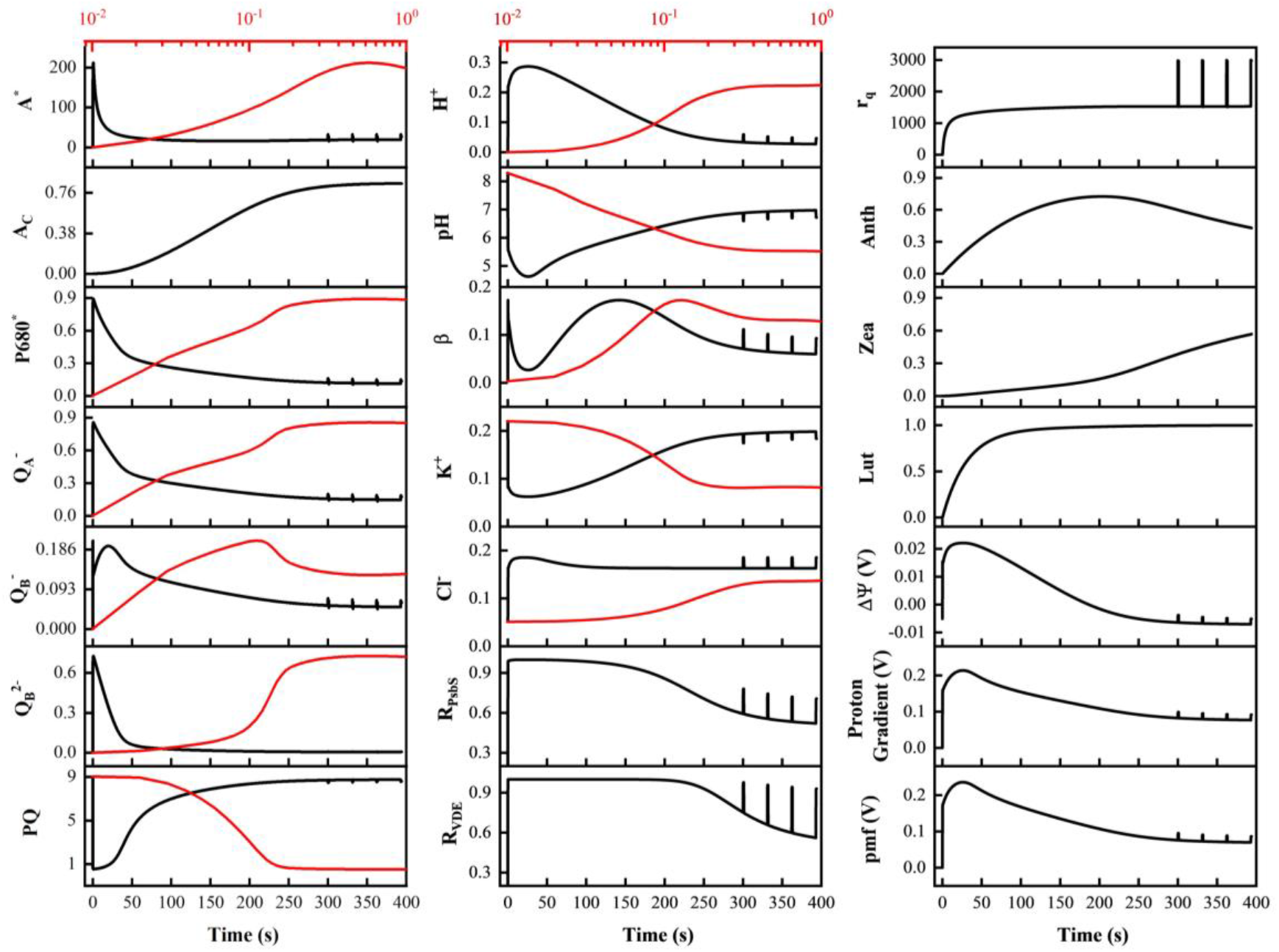
Modeled responses of 21 variables as functions of time: Activated antenna (A*), excited photosystem II primary donor (P680*), antenna-carotenoid complex (A_c_), reduced Q_A_ (Q_A_^-^), single-reduced Q_B_ (Q_B_^-^), double-reduced Q_B_ (Q_B_^2-^), plastoquinone (PQ), net proton production (H^+^), pH, buffer capacity (b), potassium (K^+^), chloride (Cl^-^), protonated PsbS (R_PsbS_), protonated VDE (R_VDE_), A* quenching rate (*r*_*q*_, mechanism 1), antheraxanthin (Anth), zeaxanthin (Zea), lutein (Lut), electric potential gradient (Δ*ψ*), ΔpH component of pmf (proton gradient effect), and proton motive force (pmf). The black curves are in the linear time scale shown at the bottom and the red curves are in log scale for the first second shown on the top. A*, P680, Q _A_^-^, Q_B_^-^, Q_B_^2-^, PQ, Anth, Zea, and Lut are fractions or probabilities.

The production of Zea and Lut leads to formation of the antenna-carotenoid complex (mechanism 2). A_c_ steadily increases and levels off to approximately 0.8, accounting for (thus “disconnecting or disabling”) a majority of the antennae in a reaction center, leading to continued and sustained declines in P680* and thus in ChlF.

The reduced energy flow from A* to P680 also slows the photochemical reactions as indicated by the low 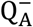and 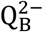. As a consequence, H^+^ decreases and pH climbs back to about 6.6, reducing R_PsbS_ to around 0.4. Because of the elevated Zea and Lut levels, nevertheless, the PsbS-mediated quenching of A* is sustained at a heightened level as shown by *r*_*q*_. Both A* quenching (at the rate *r*_*q*_) and A_c_ formation play roles in disrupting energy transfer to P680 by consuming A* or disengaging A* from P680, thus simultaneously affecting ChlF. During the pulses, *r*_*q*_ increases to consume more A* because of elevated R_PsbS_. A_c_ plays a more sustained role and results in low amplitudes of ChlF responses to the illumination pulses as will be further discussed in a later section.

The buffer capacity stays high during an illumination pulse, partially suppressing the pH decrease caused by the high proton production in the lumen by the strong illumination. The pH spikes have a minor impact on Zea and Lut, as they are regulated within the epoxide cycle and a short-time stimulation cannot lead to instantaneous changes in them. While exhibiting visually minimal responses to excitation pulses, both Zea and Lut contribute to the pulses in *r*_*q*_. Additionally, Anth also has an influence on *r*_*q*_ during light adaptation. Reduction of pH induced by pulse illumination results in a swift elevation of *r*_*q*_ because of an increase in R_PsbS_. It is worth noting that a majority of antennae are effectively decoupled from P680 by A_c_ formation, thereby establishing A_c_ as the primary determinant of the fluorescence amplitude of response to the pulse excitations.

In response to changes in H^+^, which represents the change in the proton gradient or difference across the thylakoid membrane, K^+^ and Cl^-^ vary in roughly opposite directions, and Δ*ψ* varies in a similar fashion to H^+^. Δ*ψ* is much smaller than the proton concentration potential effect ((ΔpH); thus, the resulting pmf is dominated by and approximately follows the latter, as shown in the figure.

### Two mechanisms of photoenergy regulation

As the measured data in Fig. 2 show, ChlF declined significantly in the PSMT phases after NPQ started at approximately 1 s though illumination *u* remained unchanged. This means that the *magnitude ratio* of ChlF over *u* (a measure of response to a constant stimulation, commonly referred to as “DC gain” in physical sciences included here for readers familiar with the term) is reduced by NPQ. For the sample, this ratio (or DC gain) reached a maximum of about 2.3 (2,300/1,000, peak ChlF over *u*) and became approximately 0.3 (600/2,000, peak ChlF during pulse over *u*_pulse_) during the period of pulses in illumination. The *amplitude ratio* of ChlF over *u* (a measure of change in response to change in stimulation, commonly called “AC gain” in physical sciences) is also subsequently reduced to about 0.1 (100/1,000, change in ChlF during pulse over (*u*_pulse_-*u*)) for the pulse illumination. Similar phenomena were observed from other samples as well.

While it seems intuitively possible to account for the reductions in both ratios (DC and AC gains, which respectively represent the sizes of ChlF responses to slow and fast changing illuminations) by only mechanism 1 of NPQ (quenching of A* at the rate *r*_*q*_), a careful analysis shows otherwise. The existence of mechanism 2 is evidenced in multiple ways as discussed below.

#### ChlF behavior

When only mechanism 1 is present, the model can be optimized to approximately follow either the OJIP phases (0-1 s) and the initial decline in ChlF after its maximum or the responses to the pulse excitations in light adaptation, but not for the entire experiment (Fig. 4).

**Fig. 4.**
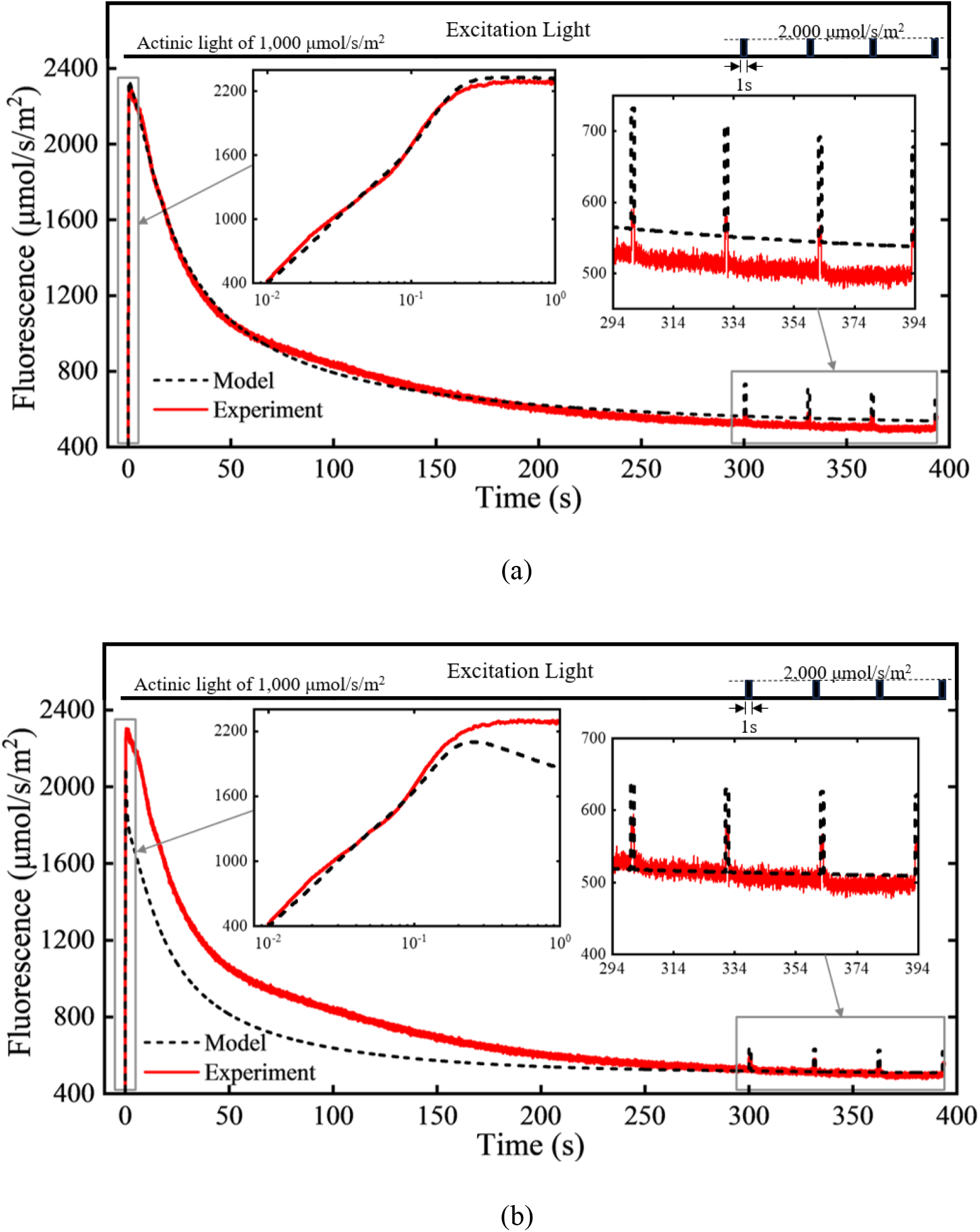
Comparison between measured and modeled ChlF for a spinach sample without A_c_ formation (mechanism 2) in the model. Insets show the OJIP phases in log scale or a magnified view of the responses to illumination pulses. (a) The model fits the measurement during the OJIP phases and ChlF decline during light adaptation, but the amplitude of responses to the illumination pulses is substantially too large, indicating insufficient A* quenching or NPQ. (b) The amplitude to the illumination pulses is closer to the measurement, but the ChlF decline starts too early and too strongly, indicating overly strong NPQ.

With only mechanism 1, an increase in illumination *u*, by the onset of actinic light for example, will begin to increase A* almost instantaneously, which would equally increase the chances for the subsequent destinations of A*, including P680 and thus ChlF. When A* quenching is appropriate for the initial responses (including maximum ChlF, thus the correct ChlF magnitude or DC gain), the amplitude of ChlF response to the pulses (or AC gain) is substantially over-predicted (Fig. 4a). For ChlF to have only a limited response to the pulse illumination (thus correct amplitude or AC gain) in the light-adapted state, A* quenching must be very strong so that fast and large NPQ is produced to divert energy from the increased A* concentration. An overly strong A* quenching will also produce too sharp and too early a reduction in ChlF (leading to an incorrect magnitude or DC gain), as shown in Fig. 4(b). In agreement with this analysis, a previous model with only mechanism 1 also showed excessively predicted qE compared with measurements(Zaks *et al*., 2012).

This structural inability of the model points to the existence of mechanism 2 (represented by A_c_ formation), which leads to not only a larger reduction in ChlF after NPQ starts but also a significant and sustained reduction in amplitude in light-adapted state. This is possible only when, during the initial light-adaptation process, some antennae are effectively disconnected from P680 by a pH-mediated mechanism. The reduction in energy transfer from antennae will result in much reduced responses to both slow and fast changes in illumination in light-adapted state. When mechanism 2 is present, the model can closely represent *both* the initial decline in ChlF and the amplitude of response to the pulse excitation as shown in Fig. 2. Because of such reductions in ChlF, previous work has suggested using new fluorescence parameters to analyze the kinetic behavior of NPQ (Bassi and Dall’Osto, 2021).

#### Trimer increase

With only mechanism 1, the number of antennae in a reaction center can be increased in the model to enhance the chance or rate of A* quenching (*r*_*q*_ in Eqn. 9) so that the model can approximate the observed ChlF amplitude of response to the illumination pulses (Fig. 4(b)). The value needed (∼430) corresponds to approximately a 67% increase in trimer LHCII content based on a monomer-to-trimer ratio of 1:6 (Wentworth, Ruban and Horton, 2004). This is in agreement with the observation that trimers were over-expressed by approximately 60% in a “no-monomer” mutant (Dall’Osto *et al*., 2017). Since monomers are believed to serve as the primary energy transfer pathway between A* and P680 (Dall’Osto *et al*., 2014; Nicol, Nawrocki and Croce, 2019), mechanism 2 should be weak or nonexistent in a no-monomer mutant. It is thus logical to expect an overexpression of trimers to compensate for the weakened or lost mechanism 2.

#### NPQ comparison

As another verification of the model and demonstration of the two mechanisms, NPQ kinetics was simulated and compared with published measurements. The wild type (WT) was simulated with both mechanisms 1 and 2 active. A variant or mutant model was created by eliminating all PSII peripheral antennae (No-LHCII) and mechanism 2. The surrounding antennae contain approximately 240 chlorophylls whereas the PSII core contains 50 (Wentworth, Ruban and Horton, 2004); consequently, the number of antennae was reduced to 50 in the mutant model. Since there were no peripheral antennae to pass energy to P680, there was no mechanism 2 to speak of. Mechanism 1 still existed to account for the PsbS-mediated quenching of the PSII core chlorophylls (Nicol, Nawrocki and Croce, 2019). A saturation illumination intensity of 5,000 μmol/m^2^/s was determined by simulation and used to simulate NPQ measurements as commonly done experimentally: F_M_ and F_M_’ were generated with a series of 1-s-wide saturation pulses and NPQ was computed as (F_M_-F_M_’)/F_M_’.

Figure 5 shows the NPQ variations with time from both the wild type (WT) and mutant (No-LHCII) models. The modeled NPQ variations are very similar, in shape and magnitude, to the actual measurements from wild-type and No-LHCII mutant Arabidopsis samples reported by Nicol et al. (Nicol, Nawrocki and Croce, 2019). Like the measurements, NPQ increases with time in a logarithmic-like fashion for both sample types. The wild type increases faster and attains a much higher level of NPQ than the mutant (Fig. 5(a)). This is evidently a synergistic result of a greater number of antennae and presence of both mechanisms. As done in Nicol et al. (Nicol, Nawrocki and Croce, 2019), the NPQ values are normalized with the respective maximum values for the two sample types (Fig. 5(b)), which reveals a greater relative rate of increase for the wild type than that for the mutant, as also shown by the published measurements. The alignment of the model with comparative measurements from wild type and mutant samples (Nicol, Nawrocki and Croce, 2019) provides further evidence for the two mechanisms.

**Fig. 5.**
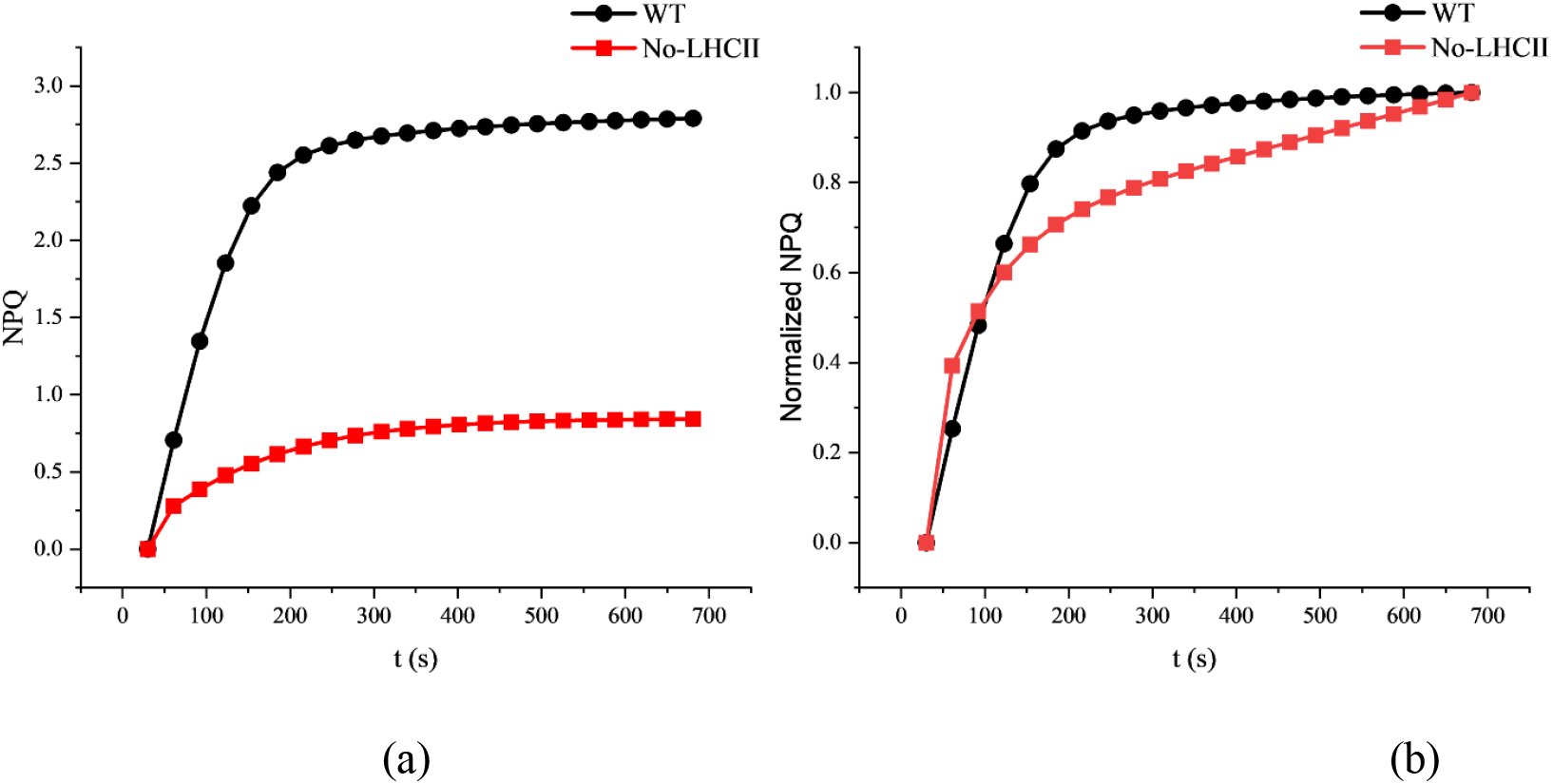
Simulated NPQ kinetics of wild type and no-trimer mutant (No-LHCII) plants. (a) NPQ computed as (F_M_-F_M_’)/F_M_’ during 1,000 μmol/m^2^/s actinic illumination. (b) NPQ normalized with respective maximum values.

#### Minimum ChlF

The minimal ChlF after light adaptation, F_0_’, is usually lower than F_0_ after dark adaptation. Bilger and Schreiber(Eni, 1967) first reported this and named it F_0_’ quenching. A similar phenomenon was also found by Bina et al. (Bína, Bouda and Litvín, 2017). The difference between F_0_’ and F_0_ indicates the existence of non-transmissibility between A* and P680 since NPQ is not expected to change significantly between the instants when F_0_’ and F_m_’ are measured (Pfündel *et al*., 2013). It is conceivable that a conformational alteration within the antennae results in a direct dissipation of energy and circumvents energy transmission to the electron transport chain, as modeled by mechanism 2.

Furthermore, the characteristics of the two mechanisms are consistent with those of the fast and the slow NPQ components reported by Bina et al. (Bína, Bouda and Litvín, 2017).

#### Concept summary

Figure 6 depicts the concept of the two mechanisms of photoenergy regulation we have observed and developed in this work. After absorbing light, antenna A becomes activated A*. A* is deactivated by passing energy to P_680_ via *k*_*A*_ or converting to heat via *k*_*NPQ*_. The excited P680* relaxes to emit fluorescence via *k*_*F*_ or drive photosynthesis via *k*_*ETR*_. Photosynthesis leads to pH changes which activate feedback actions of light adaptation. The feedback produces light-adaptation activities in two different processes via two mechanisms. Mechanism 1 leads to quenching of A* into heat as depicted in the figure by the modulation of *k*_*NPQ*_ by feedback factor *k*_*LA*1_. Mechanism 2 decreases the photoenergy transmission from activated antenna A* to P_680_ as depicted by the modulation of *k*_*A*_ by feedback factor *k*_*LA*2_.

**Fig. 6.**
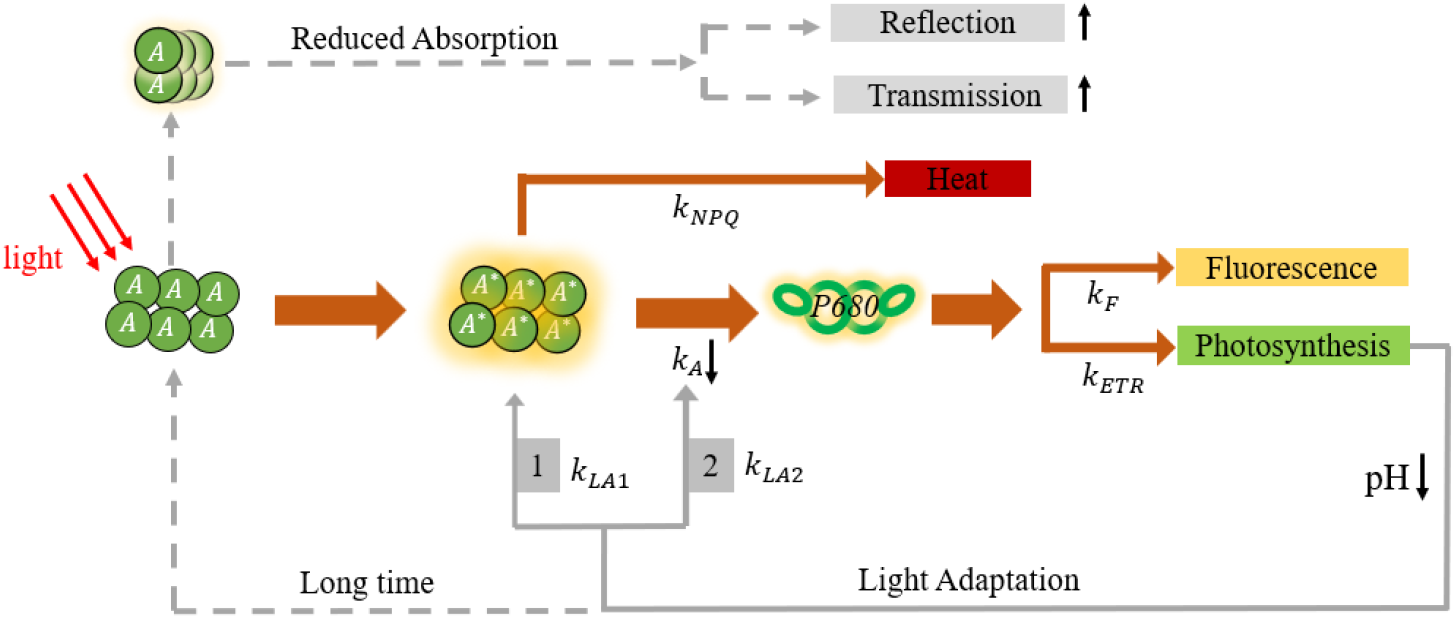
Diagram illustrating the concept of the two mechanisms of photoenergy regulation. Upon absorbing light, antenna A becomes activated A*. A* may become heat via *k*_*NPQ*_ or transfer energy to P_680_ through *k*_*A*_. P_680_* may relax to generate fluorescence via *k*_*F*_ or provides energy for photosynthesis via *k*_*ETR*_. pH reduction from photosynthesis activates feedback actions of light adaptation in two processes or mechanisms. Mechanism 1 leads to quenching of A* into heat as depicted by the modulation of *k*_*NPQ*_ by feedback factor *k*_*LA*1_. Mechanism 2 decreases the transfer of energy from antenna A* to P680 as depicted by the modulation of *k*_*A*_ by feedback factor *k*_*LA*2._

Existing knowledge on photoelectron transduction in photosystem II was synthesized in a kinetic model to allow observation of mechanisms of photoenergy regulation during light adaptation. The modeling process and experimental model validation suggest that besides pH- and PsbS-mediated nonphotochemical quenching, a disconnection between active antennae and the PSII core is necessary to account for sustained reductions of fluorescence in light-adapted conditions and to explain variations in trimer, NPQ, and minimum chlorophyll fluorescence. Model simulations provide a holistic picture of all the major kinetic variations in photosystem II. The model and analysis provide insights into the feedback mechanisms in photosystem II for regulation of photoenergy transport.

## Methods

### Experiments

Fresh *Spinacia oleracea* samples were acquired locally in Columbia, MO and used for all the experiments. A chlorophyll fluorometer (Model OS5p+, Opti-Sciences, Hudson, NH, USA) was used to provide illumination excitation and to measure ChlF. The instrument was customized by the manufacturer with expanded memory and ability to program the excitation profile used (see below). No changes were made to its light sources and sensing system.

*S. oleracea* leaf samples were dark-adapted in measurement clips for at least 30 minutes before illumination and measurement. From the dark-adapted state, a constant light of 1,000 μmol/m^2^/s was applied as actinic light to induce light adaptation. Starting at 300 s, four pulses of 2,000 μmol/m^2^/s (1,000 μmol/m^2^/s above the actinic light) were executed to generate dynamic variations in light-adapted status. Each pulse lasted for 1 s and the interval between two consecutive pulses was 30 s. The sampling interval was 1 ms in the OJIP phases (before 1 s) and 10 ms afterwards.

### Parameter estimation

The fourth-order Runge-Kutta (RK4) method^44^ was used to solve the nonlinear differential equations (Eqns. S22-36) and the model parameters were estimated by using the Levenberg–Marquardt method^45^ based on the experimentally measured ChlF. Matlab (The MathWorks, Natick, MA, USA) was used to program all the algorithms.

To evaluate the model performance against measured ChlF, the mean absolute percentage error (MAPE) and R^2^ were computed to test the agreement between model predictions and measurements. The MAPE is defined as:

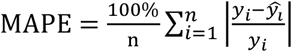

where *y* is the measured ChlF, n is the number of the data points, and ŷ is the model-predicated ChlF.

### Initial conditions

A sample is assumed to be in an initially dark-adapted steady state, from which the initial conditions can be easily determined. First, in dark-adapted state, all the photochemical reactions cease and thus ([*A*^∗^], 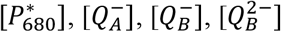, [*Anth*], [*Zea*], [*Lut*], and [*A*_*c*_] are zero. As [*H*^+^] is defined as the change from the initial dark-adapted state, it is initially zero. The initial value of [*PQ*] is *PQ*_0_ = 9.8 ^23^ and the initial *pH* in the lumen is *pH*_0_ = 7.8 ^46,47^, as mentioned previously.

In dark-adapted steady state, all the ion fluxes (Eqns. S19, S20 and S21 for *pmf*, [*K*^+^], and [*Cl*^−^], respectively) are zero. From the lumen and the stroma pH values (7.8 and 8), an initial (Δ*ψ* = -0.0169 can be determined by setting Eqn. (S17) to zero. The initial [*K*^+^] and [*Cl*^−^] values can be found to be 0.22 mol/L, and 0.0511 mol/L, respectively, by setting Eqns. (S20) and (S21) to zero and using [*K*^+^]_*str*_ = 0.15 mol/L and [*Cl*^−^]_*str*_ = 0.075 mol/L. Substituting the determined initial and known values into Eqn. (S18) yields 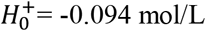. This negative value indicates that the total (both free and bond) proton concentration in the lumen is lower than that in the stroma for the initial pH values and stroma ion concentrations used, but the free proton concentration in the lumen is higher (lower pH) than that in the stroma. The former represents a negative contribution to the pmf via electrical potential (Δ*ψ* while the latter is a positive contribution via the concentration potential (ΔpH) in initially dark-adapted conditions.

## Supporting information

https://github.com/jc4q3/Matlab-Code-of-Light-adaptation-Modeling.git

## Acknowledgements

This material is based upon work supported by the US National Science Foundation under Grant No. 1903716.

## Competing interests

None declared.

## Author contributions

J.C. did measurements, developed the model, and drafted the manuscript. L.F. and Y.G. added details of the model and analyzed the data. J.T. supervised the research. All reviewed and edited the manuscript.

## Data and Code availability

Data presented in this manuscript are available and can be obtained from the corresponding author. Codes developed and used in this manuscript can be found at: https://github.com/jc4q3/Matlab-Code-of-Light-adaptation-Modeling.git

## Notes

### Competing Interest Statement

The authors have declared no competing interest.

