## Supplementary material for "Two mechanisms of photoenergy regulation revealed by kinetic behaviors of chlorophyll fluorescence during light adaptation": https://github.com/jc4q3/Matlab-Code-of-Light-adaptation-Modeling.git

### Supplementary Materials

#### Reaction Analysis and Model Development

Photochemical quenching After capturing a photon, a PSII antenna (A) is activated to become A* and the energy will transfer among the chlorophylls in the antennae. When a P680 captures the energy to become the excited form, P680*, the energy has two possible destinations. P680* may pass its energized electron, via pheophytin, to plastoquinone (PQ) molecules through plastoquinone A and B ($Q_A$ and $Q_B$) to drive subsequent photochemical reactions and receives a replacement electron originating from water(Rosenqvist and van Kooten, 2003). P680* may relax to the ground state and release chlorophyll fluorescence (ChlF). One $Q_A$ molecule carries one energized electron and a $Q_B$ can carry two. Details of electron transport for photochemical reactions were described in previous work (Guo and Tan, 2011) and the main reactions are listed below.

(i) Activation of antenna A by photon absorption

$$A + u \rightarrow A^* \quad \text{(S1)}$$

(ii) Energy transfer from activated antenna A* to P680

$$A^* + P_{680} \rightarrow A + P_{680}^* \quad \text{(S2)}$$

(iii) Relaxation of P680* to emit fluorescence

$$P_{680}^* \rightarrow P_{680} \quad \text{(S3)}$$

(iv) $Q_A$ reduction

$$P_{680}^* + Q_A \leftrightarrows Q_A^- + P_{680}^+ \quad \text{(S4)}$$

(v) $Q_B$ reduction

$$Q_A^- + Q_B \leftrightarrows Q_A + Q_B^- \quad \text{(S5)}$$

$$Q_A^- + Q_B^- \leftrightarrows Q_B^{2-} + Q_A \quad \text{(S6)}$$

(vi) PQ reduction

$$Q_B^{2-} + PQ \rightarrow Q_B + PQH_2 \quad \text{(S7)}$$

where $u$ is the illumination light intensity; $Q_A^-$ is reduced $Q_A$; $Q_B^-$ and $Q_B^{2-}$ are reduced and double-reduced $Q_B$, respectively; and $PQH_2$ is plastohydroquinone or reduced PQ. Electron transfer by pheophytin is much faster than the other steps and is thus neglected as done in previous research(Guo and Tan, 2011).

Cytochrome $b_6f$ may pump protons from the stroma to the lumen of the thylakoid and pass electrons from plastohydroquinone ($PQH_2$) to plastocyanin (PC), which is regulated by the acidification of the lumen(Kirchhoff, Li and Puthiyaveetil, 2017). The electrons are mediated by ferredoxin (Fd) and are finally transferred to NADPH through NADP+ reductase. These reactions can be summed as follows:

(vii) Electron flow from plastohydroquinone ($PQH_2$) to NADPH

$$PQH_2 + 3H_{str}^{+} + NADP^{+} \rightarrow PQ + 4H^{+} + NADPH \tag{S8}$$

where $H_{str}^{+}$ and $H^{+}$ are protons in the stroma and the lumen, respectively.

Reactions (S1-S7) are nonenzymatic and thus follow the rate laws for elementary reactions. Reaction (S8) is enzymatic as the conversion from $NADP^{+}$ to NADPH requires ferredoxin $NADP^{+}$ oxidoreductase (FNR); therefore, its rate can be described by the Michaelis-Menten equation. The net forward reaction rates for Reactions (S1-S8), $r_1$-$r_8$, can thus be expressed as follows.

$$r_1 = k_1 u(A_0 - [A^*]) \tag{S9}$$

$$r_2 = k_2(1 - [A_c])[A^*][P_{680}] \tag{S10}$$

$$r_3 = k_3[P_{680}^*] \tag{S11}$$

$$r_4 = k_4[P_{680}^*](1 - [Q_A^-]) - k_5(1 - [P_{680}^*])[Q_A^-] \tag{S12}$$

$$r_5 = k_6[Q_A^-](1 - [Q_B^-] - [Q_B^{2-}]) - k_7[Q_B^-](1 - [Q_A^-]) \tag{S13}$$

$$r_6 = k_8[Q_A^-][Q_B^-] - k_9[Q_B^{2-}](1 - [Q_A^-]) \tag{S14}$$

$$r_7 = k_{10}[Q_B^{2-}][PQ] \tag{S15}$$

$$r_8 = k_{11} pH \frac{PQ_0 - [PQ]}{k_{12} + (PQ_0 - [PQ])} \tag{S16}$$

where $k_i$ ($i$ = 1-11) are reaction rate constants; $k_{12}$ is the Michaelis constant for Reaction (S8); $A_0$ is the number of antennae in one reaction center, set to 290(Guo and Tan, 2011); $A_c$ represents the percentage of antennae disengaged from P680 for photoenergy transfer, which will be discussed later in the next subsection (Non-photochemical quenching); $PQ_0$ is the average number of PQ in a reaction center, 9.8 (Guo and Tan, 2011); $pH$ is the pH in the lumen and the initial pH is set to 7.8(Takizawa *et al.*, 2007; Tikhonov, 2013); and $[x]$ represents the number or fraction of species *x*.

Eqns. (S9-S16) are written on the basis of a single reaction center. $[Q_A]$, $[Q_A^-]$, $[Q_B]$, $[Q_B^-]$, and $[Q_B^{2-}]$ are thus concentrations in number of molecules per reaction center. $P680^{+}$ is considered to become P680 instantaneously by acquiring an electron originating from water; thus, P680 has only two possible states and $[P_{680}]$ and $[P_{680}^*]$ sum to 1. Similarly, $[Q_A]$ and $[Q_A^-]$ sum to 1; and $[Q_B]$, $[Q_B^-]$, and $[Q_B^{2-}]$ sum to 1.

Each of these concentrations may be alternatively viewed as the probability of a P680, $Q_A$ or $Q_B$ molecule in a particular state(Guo and Tan, 2011).

In the following analysis, a thylakoid is taken as the control volume of analysis. As a result, concentrations in mol/L, pH, and other conditions in the thylakoid lumen are the variables modeled, and the conditions in the stroma are used as the baseline reference or ground. The stroma conditions were assumed constant and designated with a subscript *str* as done in Reaction (S8). If a reaction rate (only $r_4$ and $r_8$) is involved in a subsequent reaction, it is multiplied by $\frac{N_{RC}}{N_A V}$ for unit conversion, where $N_{RC}$ is the number of reaction centers in a thylakoid, 2.5´10$^{6}$ (Thornber JP, Alberte RS, Hunter FA, Shiozawa JA, 1977) ; $N_A$ means Avogadro's number, 6.022´10$^{23}$; and $V$ represents the thylakoid lumen volume, 2.1´10$^{-15}$L (Beebo *et al.*, 2013).

<u>Proton motive force (pmf) and ion fluxes</u> ATP synthesis catalyzed by CF0-CF1 ATP synthase is driven by a proton motive force, which contains an electric potential difference (Dy) and a chemical concentration potential of protons (DpH) across the thylakoid membrane and is expressed as(Cruz *et al.*, 2001):

$$pmf = \Delta\psi - 2.3\frac{RT}{F}\Delta pH \qquad \text{(S17)}$$

where $R$ is the gas constant, 8.3145 J/mol/K; $T$ is absolute temperature in K; $F$ is the Faraday constant, 96,485 C/mol. Since the lumen is the control volume of analysis, D$pH = pH\text{-}pH_{str}$ with $pH_{str}$ being 8 (Werdan, Heldt and Milovancev, 1975).

The Dy or DpH component may dominate pmf during different periods. Dy is created by the electric charges across the membrane and can thus change fast, while the DpH component is generated by the chemical gradient of protons between the lumen and the stroma and its effect on pmf is much slower. Although early work suggested that DpH trumps Dy in the generation of pmf, Dy plays a first and foremost role initially during ion exchanges because of the low electrical capacitance of the thylakoid membrane, which is about 0.6 mF/cm$^2$ (Davis *et al.*, 2017). Potassium ($K^+$) and chloride ($Cl^-$) are the major ions that contribute to Dy in chloroplasts and are, therefore, used as the *representative* cation and anion, respectively, to account for the electrical effects of all ions. By Coulomb's law, Dy is:

$$\Delta\psi = \frac{e([H^+]+H_0^+ + [K^+] - [Cl^-] - [K^+]_{str} + [Cl^-]_{str})N_A V}{C_m} \qquad \text{(S18)}$$

where $e$ is the elementary charge, 1.602176634´10$^{-19}$ C; $C_m$ is electrical capacitance of the thylakoid membrane, 0.6 mF/cm$^2$ (Davis *et al.*, 2017); subscript *str* represents stroma.

The proton flux generated by pmf can be represented as:

$$r_9 = k_{13} pmf \tag{S19}$$

where $k_{13}$ is a constant representing an effective conductivity or permeability of protons across the thylakoid membrane.

The cation ($K^+$) and anion ($Cl^-$) fluxes across the thylakoid membrane from the lumen to the stroma can be similarly represented as, respectively:

$$r_{10} = k_{14}(\Delta\psi + \frac{RT}{F}\ln(\frac{[K^+]}{[K^+]_{str}})) \tag{S20}$$

$$r_{11} = k_{15}\left(-\Delta\psi + \frac{RT}{F}\ln\left(\frac{[Cl^-]}{[Cl^-]_{str}}\right)\right) \tag{S21}$$

where $k_{14}$ and $k_{15}$ are, respectively, constants accounting for the conductivity or permeability of $K^+$ and $Cl^-$ through the thylakoid membrane. The $K^+$ and $Cl^-$ channels, such as KEA3 and VCCN1, may vary in the NPQ process (Li *et al.*, 2021). These channel variations are not specifically modeled here but $k_{14}$ and $k_{15}$ were allowed to vary in the parameter estimation process. $[K^+]_{str}$ and $[Cl^-]_{str}$ are the $K^+$ and $Cl^-$ concentrations in the stroma, which are 0.15 mol/L and 0.075 mol/L, respectively (Demmig and Gimmler, 1983).

<u>Nonphotochemical quenching</u> pH is the trigger because violaxanthin de-epoxidase (VDE) and Photosystem II subunit S protein (PsbS) get protonated in low lumen pH. VDE is a very large molecule compared with proton(Pinnola and Griffiths, 2019); therefore, the proportion of protonated VDE can be expressed by the Hill equation:

$$R_{VDE} = \frac{[H^+]^{n_{P_{VDE}}}}{K_p^{n_{P_{VDE}}} + [H^+]^{n_{P_{VDE}}}} = \frac{(10^{-pH})^{n_{P_{VDE}}}}{(10^{-pKa_{VDE}})^{n_{P_{VDE}}} + (10^{-pH})^{n_{P_{VDE}}}} \tag{S22}$$

where $pKa_{VDE}$ and $n_{P_{VDE}}$ are the pKa and the Hill coefficient of VDE protonation, which equal to 6.5 and 4, respectively(Takizawa *et al.*, 2007). VDE de-epoxidates violaxanthin (Vio) to zeaxanthin (Zea) though antheraxanthin (Anth), and lutein epoxide (Lx) to lutein (Lut); i.e.,

$$\text{Violaxanthin } (Vio) \leftrightarrows \text{Antheraxanthin } (Anth) \tag{S23}$$

$$\text{Antheraxanthin } (Anth) \leftrightarrows \text{Zeaxanthin } (Zea) \tag{S24}$$

$$\text{Lutein epoxide } (Lx) \leftrightarrows \text{Lutein } (Lut) \tag{S25}$$

Based on the Michaelis-Menten enzymatic kinetics, the net forward reaction rates of Reactions S23-S25, $r_{12}$ to $r_{14}$, are:

$$r_{12} = \frac{V_{maxVA} R_{VDE}(1-[Anth]-[Zea])}{k_{mVA}+1-[Anth]-[Zea]} - \frac{V_{maxAV} R_{ZE}[Anth]}{k_{mAV}+[Anth]} \quad \text{(S26)}$$

$$r_{13} = \frac{V_{maxAZ} R_{VDE}[Anth]}{k_{mAZ}+[Anth]} - \frac{V_{maxZA} R_{ZE}[Zea]}{k_{mZA}+[Zea]} \quad \text{(S27)}$$

$$r_{14} = \frac{V_{maxL} R_{VDE}(1-[Lut])}{k_{mL}+(1-[Lut])} \quad \text{(S28)}$$

Early studies suggested that PsbS , discovered in 1981(Berthold, Babcock and Yocum, 1981), was the site of NPQ in photosynthesis and was able to bind with chlorophylls and carotenoids; however, it was later shown that PsbS would not directly bind with pigments because of its extreme hydrophobicity(Dominici *et al.*, 2002; Barbara Demmig-Adams, Gyozo Garab, William Adams III, 2014). This property means that it reacts with the pigment-binding protein in the antennae instead of quenching energy by itself(Pinnola and Griffiths, 2019). The process of PsbS protonation meets the requirements of the Hill equation and thus the proportion of protonated PsbS can be computed as:

$$R_{PsbS} = \frac{(10^{-pH})^{n_{P_{PsbS}}}}{(10^{-pKa_{PsbS}})^{n_{P_{PsbS}}}+(10^{-pH})^{n_{P_{PsbS}}}} \quad \text{(S29)}$$

where $pKa_{PsbS}$ and $n_{P_{PsbS}}$ are the pKa and the Hill coefficient of PsbS protonation, which are set to be 6.5 and 1.36, respectively(Johnson and Ruban, 2011).

The rate of quenching of activated antenna A* by carotenoids can be expressed as:

$$r_{15} = R_{PsbS}(k_{16}[Anth] + k_{17}[Zea] + k_{18}[Lut])[A^*] \quad \text{(S30)}$$

where $k_{16-18}$ are the reaction rates for Lut, Zea and Anth, respectively, which are symbolized with $k_{A,Z,L}$ in the main text; and $r_{15}$ is $r_q$ in the main text.

Zea and Lut will form complex $A_c$ with antenna and the complexed antennae are conformationally changed and thus unable to pass energy to P680, as detailed in the main text. The rates of complex formation are:

$$r_{16} = \frac{V_{max1} R_{PsbS}[Zea](A_{c0}-[A_c])}{k_{mA1}+[Zea]+A_{c0}-[A_c]} \quad \text{(S31)}$$

$$r_{17} = \frac{V_{max2} R_{PsbS}[Lut](A_{c0}-[A_c])}{k_{mA1}+[Lut]+A_{c0}-[A_c]} \quad \text{(S32)}$$

where $r_{16,17}$ are $r_{A_{Zea},A_{Lut}}$ in the main text; $V_{max1,2}$ and $k_{mA1,2}$ represent the maximum rate and Michaelis constant of $A_c$ formation from Zea and Lut, respectively; $[A_c]$ represents the proportion of conformationally changed antennae; $A_{c0}$ is the maximum proportion, set to 0.85(Wentworth, Ruban and Horton, 2004).

#### Model Structure

A kinetic model can be created based on the equations and reactions discussed in the preceding subsections. Fourteen variables: $A^*$, $P680^*$, $Q_A^-$, $Q_B^-$, $Q_B^{2-}$, $PQ$, $H^+$ $pH$, $K^+$, $Cl^-$, $Anth$, $Zea$, $Lut$, and $A_c$ can be represented by the following differential equations according to reactions and equations discussed above:

$$\frac{d[A^*]}{dt} = r_1 - r_2 - r_{15} \quad \text{(S33)}$$

$$\frac{d[P680^*]}{dt} = r_2 - r_3 - r_4 \quad \text{(S34)}$$

$$\frac{d[Q_A^-]}{dt} = r_4 - r_5 - r_6 \quad \text{(S35)}$$

$$\frac{d[Q_B^-]}{dt} = r_5 - r_6 \quad \text{(S36)}$$

$$\frac{d[Q_B^{2-}]}{dt} = r_6 - r_7 \quad \text{(S37)}$$

$$\frac{d[PQ]}{dt} = -r_7 + r_8 \quad \text{(S38)}$$

$$\frac{d[H^+]}{dt} = \frac{N_{RC}}{N_A V}\left(r_{4f} + 4r_8\right) - r_9 \quad \text{(S39)}$$

$$\frac{dpH}{dt} = -\frac{1}{\beta}\left(\frac{N_{RC}}{N_A V}\left(r_{4f} + 4r_8\right) - r_9\right) \quad \text{(S40)}$$

$$\frac{d[K^+]}{dt} = -r_{10} \quad \text{(S41)}$$

$$\frac{d[Cl^-]}{dt} = -r_{11} \quad \text{(S42)}$$

$$\frac{d[Anth]}{dt} = r_{12} - r_{13} \quad \text{(S43)}$$

$$\frac{d[Zea]}{dt} = r_{13} - r_{16} \quad \text{(S44)}$$

$$\frac{d[Lut]}{dt} = r_{14} - r_{17} \quad \text{(S45)}$$

$$\frac{d[A_c]}{dt} = r_{16} + r_{17} \quad \text{(S46)}$$

where $r_{4f}$ is the forward reaction rate of Reaction (S4) and all other symbols have been defined previously under the rate equations (Eqns. (S9-S16)). Only the forward reaction rate of Reaction (S4) is used here because it is the proton production rate from splitting water and the backward reaction does not lead to consumption of $H^+$ (Renger, 2011).

The ChlF from one reaction center is $k_3[P_{680}^*]$ (Reaction S3, Eqn. S11). The total ChlF intensity measured in an observation area can be written as:

$$ChlF = Gk_3[P_{680}^*] + F_0 \qquad \text{(S47)}$$

where $G$ is an overall gain factor accounting for the instrumentation gain and the number of reaction centers in the observation area. $F_0$ represents ChlF from other mechanisms, such as Photosystem I, which is often assumed to be small and constant.

**Initial conditions**

A sample is assumed to be in an initially dark-adapted steady state, from which the initial conditions can be easily determined. First, in dark-adapted state, all the photochemical reactions cease and thus ($[A^*]$, $[P_{680}^*]$, $[Q_A^-]$, $[Q_B^-]$, $[Q_B^{2-}]$, $[Anth]$, $[Zea]$, $[Lut]$, and $[A_c]$ are zero. As $[H^+]$ is defined as the change from the initial dark-adapted state, it is initially zero. The initial value of $[PQ]$ is $PQ_0$ = 9.8 (Guo and Tan, 2011) and the initial $pH$ in the lumen is $pH_0$ = 7.8 (Takizawa *et al.*, 2007; Tikhonov, 2013), as mentioned previously.

In dark-adapted steady state, all the ion fluxes (Eqns. S19, S20 and S21 for *pmf*, $[K^+]$, and $[Cl^-]$, respectively) are zero. From the lumen and the stroma pH values (7.8 and 8), an initial $\Delta\psi$ = -0.0169 can be determined by setting Eqn. (S17) to zero. The initial $[K^+]$ and $[Cl^-]$ values can be found to be 0.22 mol/L, and 0.0511 mol/L, respectively, by setting Eqns. (S20) and (S21) to zero and using $[K^+]_{str}$ = 0.15 mol/L and $[Cl^-]_{str}$ = 0.075 mol/L. Substituting the determined initial and known values into Eqn. (S18) yields $H_0^+$= -0.094 mol/L. This negative value indicates that the total (both free and bond) proton concentration in the lumen is lower than that in the stroma for the initial pH values and stroma ion concentrations used, but the free proton concentration in the lumen is higher (lower pH) than that in the stroma. The former represents a negative contribution to the pmf via electrical potential $\Delta\psi$ while the latter is a positive contribution via the concentration potential (ΔpH) in initially dark-adapted conditions.

**Additional Samples for Model Validation**

The model was compared against measured ChlF from additional spinach samples. Fig. S1. compares model predictions with measurements from two more samples. Both show good agreement. Table S1 lists the parameter values used for the three samples.

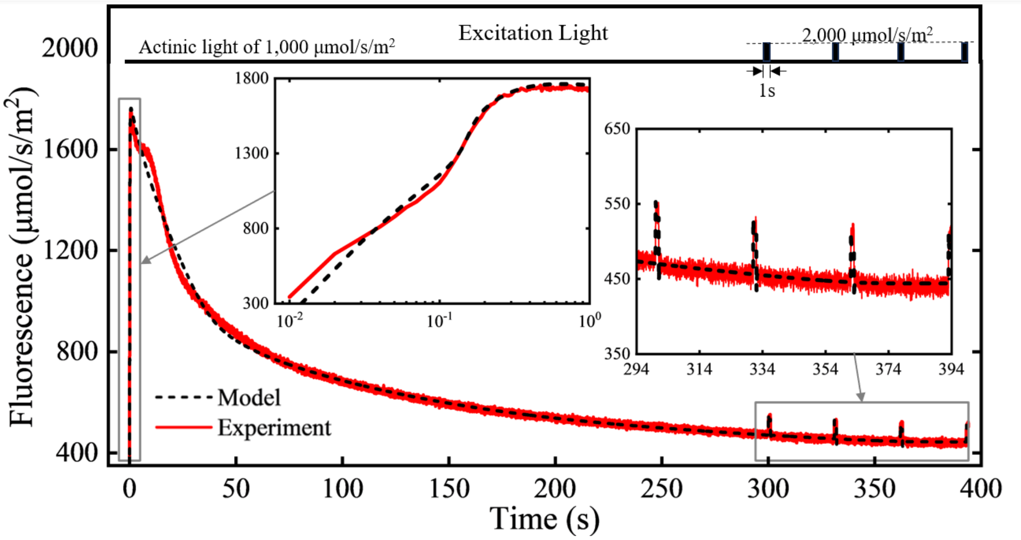


(a)

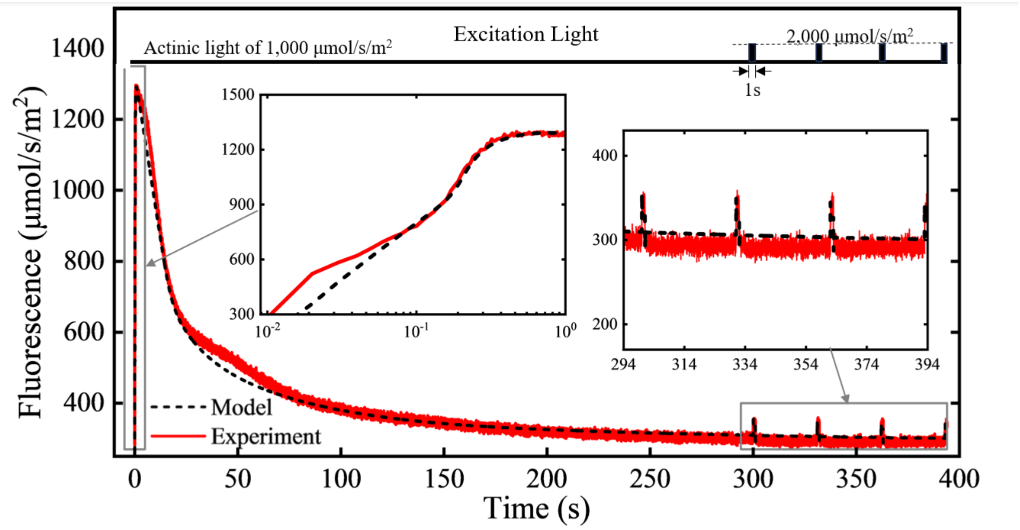


(b)

**Fig. S1 | Mode predictions compared with measured ChlF for two additional spinach samples**. The sample was an initially dark-adapted spinach leaf in 22°C. The illumination was a 1,000 mmol/s/m$^2$ constant actinic light starting at time 0 with 1-s wide pulses of 1,000 mmol/s/m$^2$ above the constant light. The light source was the built-in white LED of an OS5p+ Chlorophyll Fluorometer (Opti-Sciences, Hudson, NH, USA). One inset shows the OJIP phases in the first second in log scale and another shows a magnified view of the pulse responses.

**Table S1** Model parameter values.

| | Fig. 1 | Fig. S1.(a) | Fig. S1.(b) | | Fig. 1 | Fig. S1.(a) | Fig. S1.(b) |
|---|---|---|---|---|---|---|---|
| $k_1$ | 0.005 | 0.005 | 0.004 | $k_{16}$ | 49.729 | 18.628 | 0.128 |
| $k_2$ | 11.737 | 11.737 | 11.737 | $k_{17}$ | 93.920 | 19.155 | 753.279 |
| $k_3$ | 33.305 | 37.064 | 40.612 | $k_{18}$ | 2.127 | 34.233 | 107.648 |
| $k_4$ | 2,999.648 | 4,132.058 | 3,783.588 | $V_{maxVA}$ | 0.017 | 0.007 | 0.007 |
| $k_5$ | 4,468.241 | 2,383.438 | 1,647.293 | $k_{mVA}$ | 0.093 | 0.093 | 0.093 |
| $k_6$ | 3,137.892 | 2,295.037 | 1,984.354 | $V_{maxAZ}$ | 8.269 | 1.626 | 10.840 |
| $k_7$ | 4,682.323 | 4,798.054 | 4,786.446 | $k_{mAZ}$ | 1,299.509 | 1,299.509 | 1,299.509 |
| $k_8$ | 4,842.045 | 2,991.689 | 3,093.145 | $V_{maxL}$ | 4.538 | 5.272 | 5.196 |
| $k_9$ | 32.998 | 1,390.614 | 952.069 | $k_{mL}$ | 137.030 | 137.0297 | 137.030 |
| $k_{10}$ | 98.035 | 799.410 | 635.686 | $V_{max1}$ | 0.097 | 2.824 | 0.239 |
| $k_{11}$ | 351.810 | 492.116 | 557.538 | $k_{mA1}$ | 0.524 | 0.0439 | 1.388 |
| $k_{12}$ | 5.241 | 5.241 | 5.241 | $V_{max2}$ | 0.003 | 0.040 | 0.015 |
| $k_{13}$ | 10.188 | 13.188 | 12.188 | $k_{mA2}$ | 0.862 | 5.373 | 2.820 |
| $k_{14}$ | 162.216 | 162.216 | 162.216 | $G$ | 261.660 | 187.139 | 128.117 |
| $k_{15}$ | 46.633 | 46.633 | 46.633 | $F_0$ | 218.000 | 218.000 | 178.000 |

#### DCMU Experiment and Model Simulations

ChlF was measured from DCMU-treated spinach samples with two levels (40% and 100%) of the modulation light of the OS5p+ Chlorophyll Fluorometer as excitation (without actinic lighting). $k_4$ and $k_5$ were set to zero in the model to simulate the disruption of electron transport by DCMU. The model was able to follow the measured ChlF (Fig. S2).

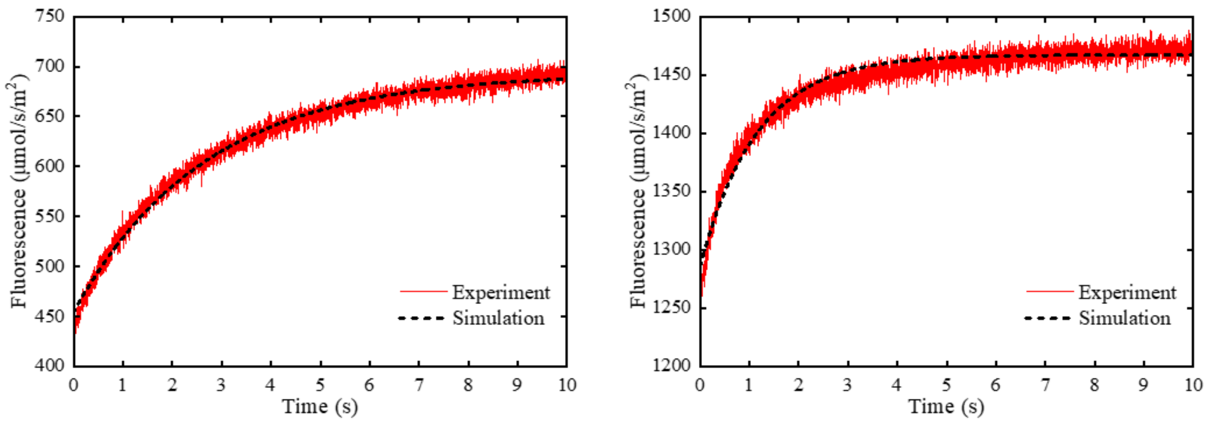


**Fig. S3 | Measured ChlF from DCMU-treated spinach samples and model simulations.** Excitation was 40% (left) or 100% (right) of the modulation light of the OS5p+ Chlorophyll Fluorometer. $k_4$ and $k_5$ were set to zero in the model to simulate disruption of electron transport by DCMU.
